# Competitive environment predicts weaponry in an intertidal sea anemone

**DOI:** 10.64898/2026.05.17.725755

**Authors:** Sriram V. Ramamurthy, Jacob G. Stinnett, Carter S. Kaulback, Ashley T. Berry, Todd H. Oakley

## Abstract

Animal weapons are ecologically important traits that mediate contests over limiting resources and can strongly influence survival and reproduction. Weapon traits often exhibit substantial intraspecific morphological diversity, raising questions about the ecological drivers of this variation. Acrorhagi are weapons produced by sea anemones that are used in intraspecific territorial encounters. Although acrorhagial morphology varies widely within species, patterns of intraspecific variation remain poorly characterized, and the extent to which such variation reflects differences in local intraspecific competition is unclear. Here, we conduct morphometric analyses to characterize within-population variation and allometry in acrorhagial traits of the solitary anemone *Anthopleura sola*. We show that these traits covary with habitats differing in conspecific density. The number of acrorhagi scaled positively with body size, and individuals occupying a high-density habitat tended to possess more acrorhagi than did similar sized individuals from a low-density habitat. In addition, anemones from high-density habitats exhibited longer acrorhagial cnidae, a pattern that was not explained by differences in body size or acrorhagial density. Together, these results suggest that competitive context influences weapon-related traits at multiple levels of biological organization, potentially via phenotypic plasticity or selective processes. More broadly, our findings highlight how fine-scale ecological variation may contribute to the maintenance of trait diversity within and across species.

## Introduction

Across diverse taxa, animal weapons are used in both intra- and interspecific conflict, influencing access to space, resources, and mating opportunities (Emlen, 2008; Lane, 2018; Miller *et al*., 2026). Because the outcome of agonistic encounters can depend strongly on weapon form and performance, the nature of an individual’s weaponry may be expected to have direct consequences for survival and reproductive success. Like many other traits (e.g., Gamelon *et al*., 2025), weapon-related traits frequently exhibit morphological variation, even among individuals of the same species (Goczal *et al*., 2019; Emberts *et al*., 2021). Such variation may reflect local ecological conditions, one aspect of which may be the social environment. Weapon morphology may be expected to exhibit plastic or selective responses under conditions that increase the probability of encountering rivals, such as high local density. However, despite this expectation, relatively little is known about how weapon traits vary as a function of social context within natural populations. Some studies have provided evidence for density-dependent responses in sexually selected competitive traits (for a review, see Knell, 2009). For example, in European earwigs, populations at higher densities exhibit a greater proportion of long-forceps males (Tomkins and Brown, 2004), and across chernetid pseudoscorpions, density is positively associated with the degree of sexual dimorphism in claw size (Zeh, 1987). Yet for other systems, weapon investment may be reduced under high density conditions (Knell, 2009). Overall, it is still unclear to what extent intraspecific variation in weapon morphology reflects selective or plastic responses to local social environments.

Acrorhagi are weapons used by actiniid sea anemones (Actiniaria: Actiniidae) exclusively in territorial contests with other conspecific and heterospecific anemones. Located at the distal edge of the body column (the margin), acrorhagi are bulbous protuberances containing dense batteries of cnidae, or stinging organelles (Daly, 2003). Detailed information about acrorhagial anatomy is provided in the Supplementary Material. Upon contact with another individual, anemones inflate their acrorhagi and engage in a series of complex agonistic behaviors (Fig. 1B, 2B) (Francis, 1973a; Bigger, 1980; Sebens, 1984). The acrorhagial ectoderm peels away and adheres to the victim, causing tissue necrosis via venom injected by the associated cnidae. Contest losers typically lean or move away, although in severe cases they may perish (Francis, 1973a,b).

**Fig. 1:**
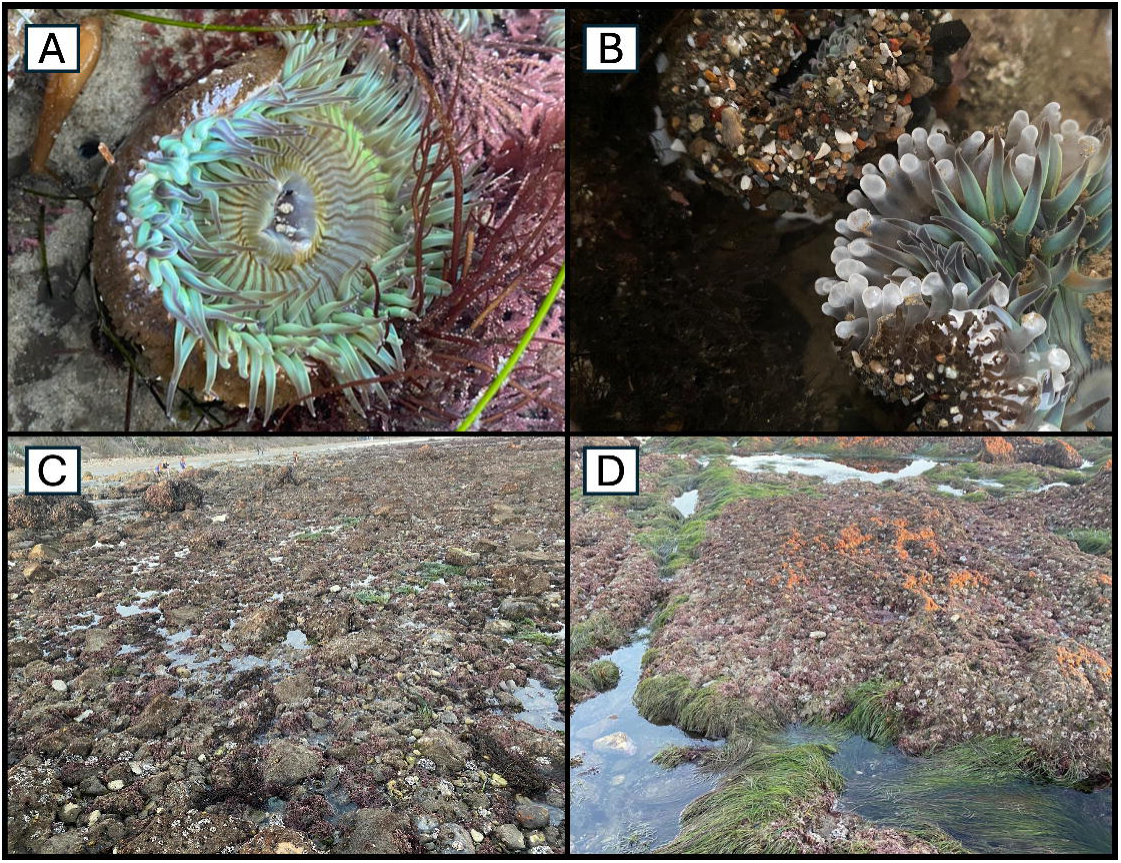
(A) *A. sola in situ*, with uninflated acrorhagi visible. (B) Agonistic episode between two individuals; the anemone on the right has inflated the acrorhagi. (C and D) cobble and bench habitats, respectively, Coal Oil Point, Santa Barbara, California.

Across acrorhagi-bearing sea anemones, acrorhagial size, shape, arrangement, and density can vary considerably within populations (e.g., Francis, 1976; Donoghue *et al*., 1985; Shick, 1991; Ramamurthy, 2024; Durán-Fuentes *et al*., 2025; V-Boada *et al*., 2026). Moreover, acrorhagi may be transient, produced in response to repeated allogeneic contact (Ayre and Grosberg, 2005), or degenerated in the prolonged absence of such stimulation (pers. obs., SVR); however, the frequency, capacity, and natural occurrence of such plastic processes are poorly studied. Variability can additionally manifest at the cellular level of the cnidom (the complement of cnidae in a given tissue or individual). Cnidae can differ in size across individuals, including those present within acrorhagial tissues (e.g., Fautin, 1988; Watts and Thorpe, 1998; Kramer and Francis, 2004; Francis, 2004). These differences can be predictable or not; for example, cnidae scale with anemone body size in two *Anthopleura* species (Francis, 2004), yet no such scaling relationships have been recovered in other actinians (Kaposi *et al*., 2023).

The factors driving variation in acrorhagial morphology within species remain unclear. Some evidence suggests that intraspecific differences in acrorhagial traits may, in part, reflect the local competitive regime. For instance, the intertidal anemone *Anthopleura elegantissima* (Brandt, 1835) forms large clonal aggregations in which individuals at inter-clonal borders bear more acrorhagi than those within clonal centers (Francis, 1976; Ayre and Grosberg, 2005), possibly due to increased encounters with nonclonemates. Similarly, individuals of *Oulactis coliumensis* (Riemann-Zürneck and Gallardo, 1990) possess larger acrorhagi in denser populations (Spano *et al*., 2022). Yet, overall, it is unclear to what extent intraspecific variation in acrorhagial traits reflects underlying ecological differences.

Gross acrorhagial characteristics and their complement of cnidae are readily quantifiable, and, as with other animal weapons, both individual-scale and cellular-scale traits have been linked to contest success (Ayre and Grosberg, 1996; Rudin and Briffa, 2011; Palaoro and Peixoto, 2022). Unlike many animal weapons, however, acrorhagi are employed not in intrasexual competition for mates, but in territorial interactions with both conspecific and heterospecific competitors (Francis, 1973a; Emlen, 2008). This broader ecological role highlights acrorhagi-bearing sea anemones as a promising model system for studying trait–environment relationships.

Here, using specimens that were collected from the spring and summer months of 2024–2025, we characterize intraspecific variation and allometry in acrorhagi of a solitary eastern Pacific intertidal sea anemone, *Anthopleura sola* (Pearse and Francis, 2000) (Fig. 1A). In particular, we examine whether acrorhagial traits covary with natural, fine-scale spatial variation in local competitive environments. We assessed acrorhagial morphology at two levels of biological organization in individuals occupying multiple habitats, including two habitat types that differ in local conspecific density. We also characterized allometric relationships describing variation in acrorhagial traits; including the total number of acrorhagi possessed by each anemone, as well as the lengths of the cnidae present within acrorhagial tissues. Even though multiple different cnida types are present in the acrorhagi of *A. sola*, we specifically chose to examine the spirocyst, as this cnida type is easily identifiable and is synapomorphic for Hexacorallia (Francis, 2004), allowing the possibility of future inter-structural, intraspecific, and interspecific comparative analyses while mitigating phylogenetic confounds. We find that the number of acrorhagi exhibits negative allometry with respect to anemone body mass. Also, we show evidence that anemones from a higher-density habitat tend to possess more acrorhagi and longer acrorhagial spirocysts on average. Together, these results indicate that weapon-related traits can covary among individuals experiencing different local environmental contexts, even across small spatial scales.

## Materials and Methods

### Collection and processing

Individuals of *A. sola* across a range of sizes were collected from rocky intertidal habitats at Coal Oil Point (Santa Barbara, CA) in April 2024, May 2024, and May 2025. The collection areas of interest include a ‘cobble’ habitat (Fig. 1C), where larger anemones occur interspersed amongst small cobbles, and a ‘bench’ habitat (Fig. 1D), where smaller anemones occur over sections of a relatively flat rugose substrate. Anemones in bench habitats can occur at substantially higher local densities than those in cobble habitats (typically on the order of two to five times higher (unpublished data SVR)). The habitat areas are close in proximity (tens of meters) and are similar in tidal height. Additional anemones were collected from Rincon Point (Ventura, CA) in September 2025. Anemones were lifted off the rocky substrate by working a spatula under the pedal disk and held in flow-through aquaria for up to ∼three months before processing. To account for mass loss during aquarium holding, we applied a correction based on empirically measured weight-loss patterns (following Sebens, 1981; see Supplementary Material for details). No correction was applied to anemones from Rincon Point due to their brief holding period (approximately one week).

Acrorhagi were counted for each anemone following relaxation with menthol and/or magnesium chloride. We then counted acrorhagi either post-fixation in 10% seawater-formalin, or pre-fixation, where the limp anemone was pinned to a wax tray and the flaccid tentacles folded back to expose the margin. We took tissue samples from each anemone, including one to three endocoelic acrorhagi (in areas where the structures were well-developed), preserving them in 10% seawater-formalin. Wet weights of anemones were taken after cleaning the individuals of debris and firmly blotting with paper towels to remove excess fluid from the coelenteron.

To measure spirocysts, tissue-smears were prepared from formalin-preserved acrorhagial tissue from arbitrarily selected anemones spanning a range of sizes. Spirocysts were photographed under 1000X magnification. 20–30 capsules in intact condition were targeted for measurement per anemone. As spirocyst capsules can exhibit different degrees of curvature, the length of each capsule was measured using a spline-fitted segmented line using *ImageJ* (Schneider *et al*., 2012) tracing along the median curve of each spirocyst (Fig. 2D). We quantified spirocyst length only; cnida length generally exhibits greater biologically meaningful variation than width and is therefore the more informative axis for measuring cnida size (Francis, 2004). Lastly, the internal anatomy of some specimens was examined, with regard to the number of mesenterial pairs (which will be equal to the number of endocoels).

**Fig. 2:**
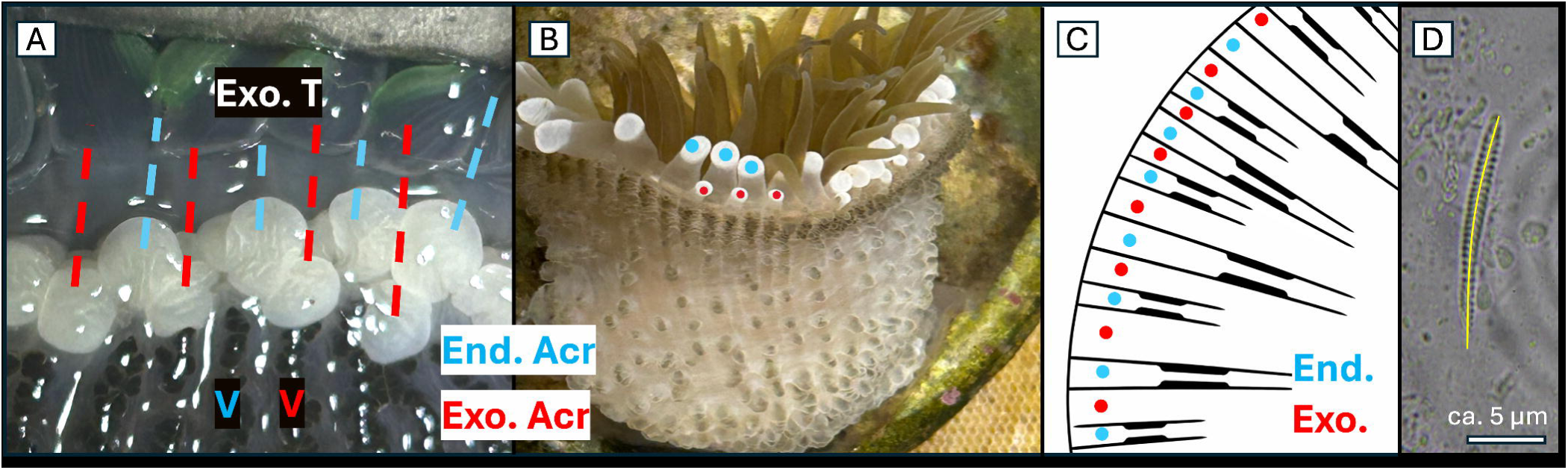
Endocoelic and exocoelic acrorhagi in *A. sola*. (A and B) relationships between positions of exocoelic outer tentacles, verrucae, and the endo- and exocoelic acrorhagi. End. Acr: Endocoelic acrorhagi. Exo. Acr: Exocoelic acrorhagi. Exo. T: Exocoelic (outer) tentacles. V: Verrucae (colored in the same respective manner as the endo- and exocoelic acrorhagi). (A) the dashed lines show the alignment of endo- and exocoelic acrorhagi with and between the exocoelic tentacles. (B) inflated endo- and exocoelic acrorhagi are denoted by appropriately colored dots. Note the smaller size of the exocoelic acrorhagi. (C) schematic representation of mesenterial arrangements in actiniid sea anemones, showing the positions of the endocoels (End.) and exocoels (Exo.) as appropriately colored dots in relation to mesenterial pairs in a cross section. (D) acrorhagial spirocyst with an example of how length was measured (yellow line).

### Statistical analysis

Analyses of acrorhagial quantity and density were conducted using linear models with logarithmic scales. As one individual possessed zero acrorhagi, we compared conventional log–log linear models (excluding the zero) with negative binomial generalized linear models (including all individuals). All analyses of spirocyst length were conducted using linear mixed-effects models, with anemone identity as a random intercept to account for non-independence of measurements, since multiple spirocysts were measured per anemone. In some instances, pairs of candidate models were ranked using Akaike’s Information Criterion (AIC). For models within ΔAIC ≤ 2, the simpler model was selected. Log-transformed predictors were mean-centered prior to analysis. Statistical analyses were performed in R (v4.5.0; R Core Team, 2025).

## Results

### Acrorhagial anatomy and morphology

Variation in acrorhagial number and organization occurs in accordance with an underlying structural (anatomical) framework. Acrorhagi are positioned at the distal ends of longitudinal rows of verrucae, with a primary cycle typically associated with endocoelic rows. In smaller or less developed individuals, acrorhagi may occur on alternating endocoels (e.g., Fig. 3B), whereas a more typical condition is a complete endocoelic cycle (e.g., Fig. 3C). With increasing size or developmental stage, a secondary cycle of exocoelic acrorhagi may arise, though this cycle is often incomplete (e.g., Fig. 3D) and composed of smaller structures positioned just below the endocoelic rows, producing a characteristic double crown (e.g., Fig. 3E–F, cf. Fig 2B). In more advanced stages, both endocoelic and exocoelic cycles may be fully developed and similar in size. Several individuals sectioned at the mid-column region possessed approximately 90–100 pairs of mesenteries, whereas one individual possessed fewer pairs (73; Suppl. Table 1).

**Fig. 3:**
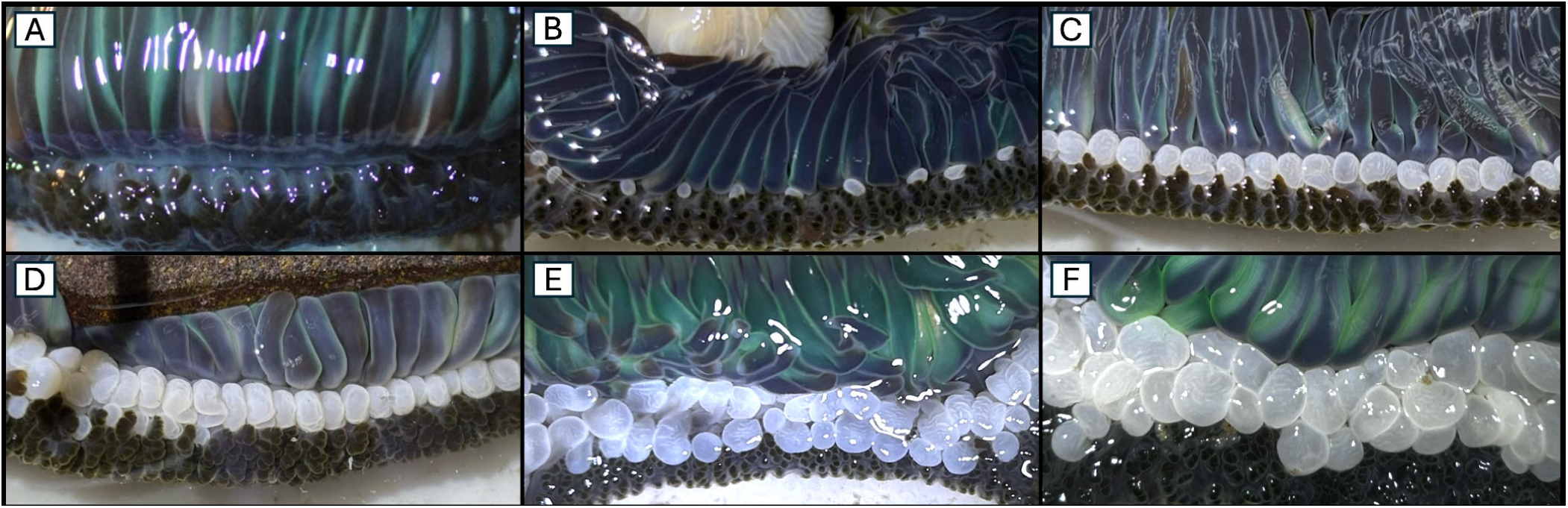
Examples of differential acrorhagial manifestation in *A. sola*. The marginal region of individuals are shown with the tentacles folded back. (A) acrorhagi absent. (B) smooth, ovoid acrorhagi present on every other endocoel. (C) ovoid and folded acrorhagi present on every endocoel. (D) ovoid and folded acrorhagi present on every endocoel, with some present on exocoels. (E) ovoid and folded acrorhagi present on every endocoel and exocoel. (F) well-developed ovoid and folded acrorhagi present on every endocoel and exocoel, and possibly some in a third cycle.

### Acrorhagial scaling

The number of acrorhagi scaled positively with anemone body mass (Fig. 4). Across all habitats, a log–log model indicated that acrorhagial number increased with log body weight (β = 0.344 ± 0.037 SE, *t* = 9.21, *p* < 0.001), consistent with negative allometry. A negative binomial GLM with a log link yielded a similar scaling exponent (β = 0.329 ± 0.038 SE, *z* = 8.57, *p* < 0.001). When analysis was restricted to bench and cobble habitats and habitat was included as an additive effect (Bench as the reference category) acrorhagial number continued to scale positively with body size. After accounting for body size, cobble anemones tended to possess fewer acrorhagi than bench anemones; statistical support for this difference varied across model formulations (log-log model: β = −0.241 ± 0.107 SE, *t* = −2.25, *p* = 0.028; negative binomial GLM (log-link): β = −0.207 ± 0.109 SE, z = −1.90, *p* = 0.058).

**Fig. 4:**
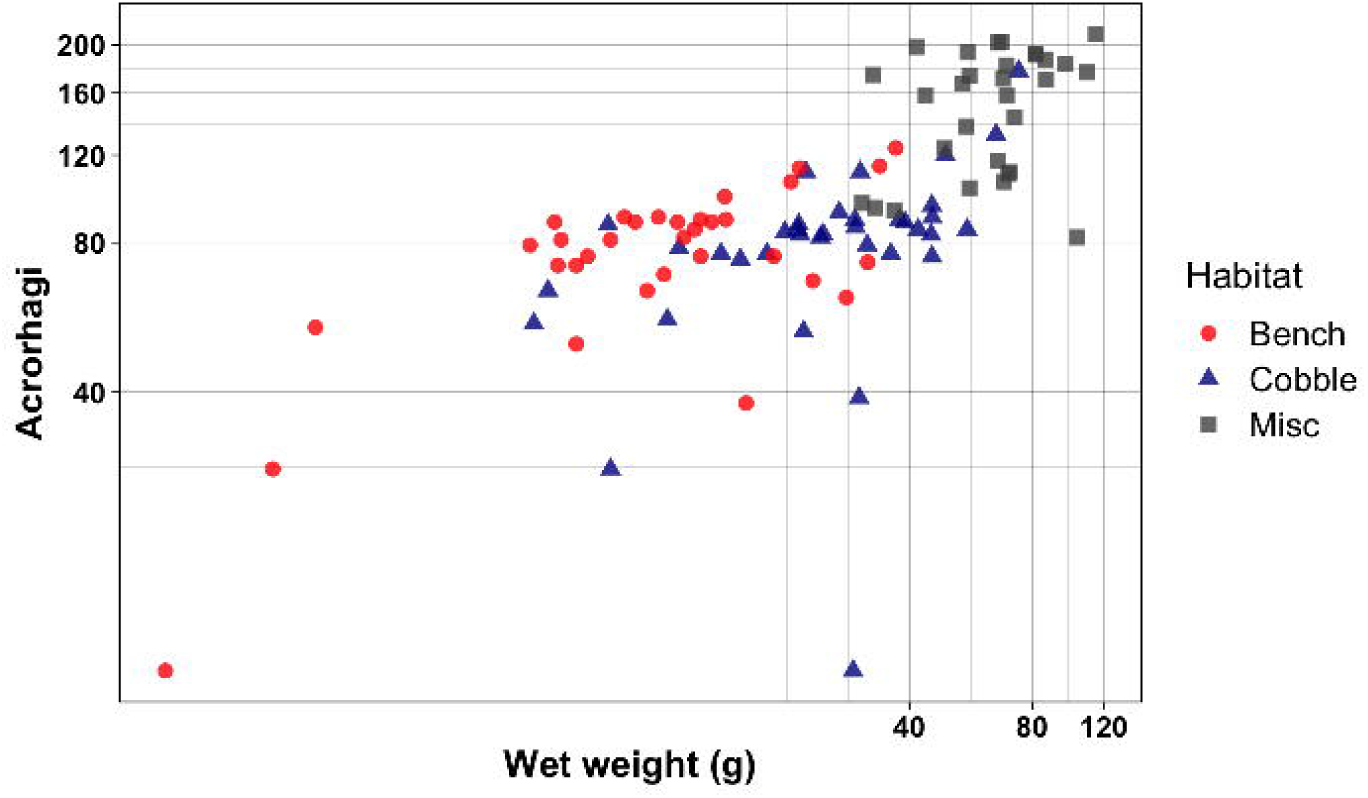
Relationship between acrorhagi quantity and wet weight (g) between habitats. Both axes are shown on logarithmic scales. One animal with 0 acrorhagi (48.7 g) is excluded from this graph. Habitats of interest include the bench (red circles) and cobble (blue triangles) habitats (Coal Oil Point, Santa Barbara), anemones collected from Rincon Point, Ventura, are also shown (“Misc”) (grey squares). N= 37, N= 33, N= 30

### Spirocyst scaling

Acrorhagial spirocyst length differed significantly between habitat types (Fig. 5). Anemones from cobble habitats possessed significantly shorter spirocysts than did those from bench habitats (estimate for cobble: −2.59 ± 0.75 SE, *t* = −3.44, *p* = 0.002). This difference corresponds to an average reduction of approximately 2.6 µm in spirocyst length for cobble anemones relative to bench anemones.

**Fig. 5:**
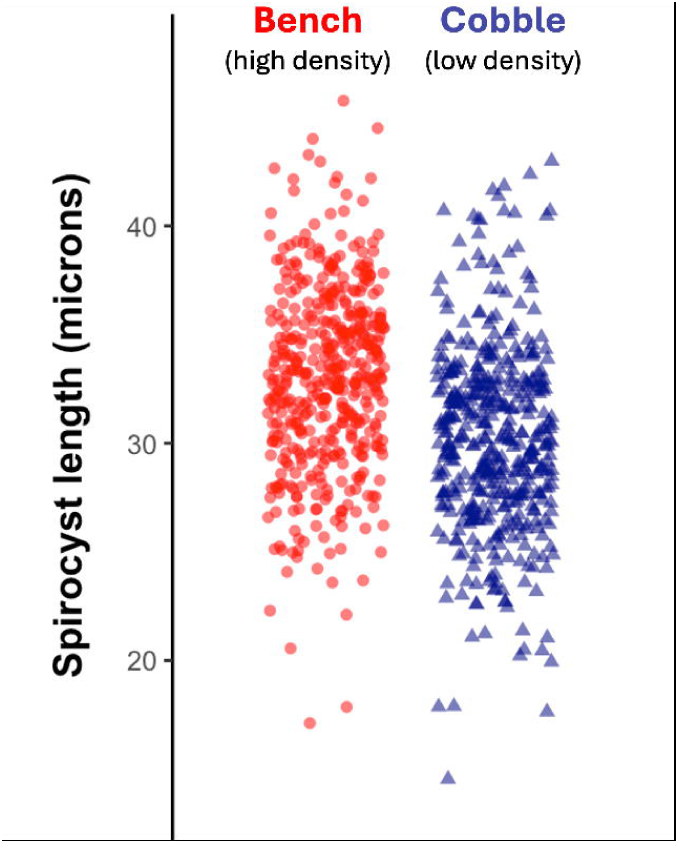
Spirocyst lengths, pooled across individuals, from anemones across bench (red circles) and cobble (blue triangles) habitats. N= 19, N= 19, N= 884

Spirocyst length differed significantly between habitat types after accounting for body size (Fig. 6). In a linear mixed-effects model including habitat and centered log body weight as an additive effect, habitat had a significant effect on spirocyst length. Cobble anemones exhibited spirocysts approximately 2.91 µm shorter than bench anemones (β = −2.91 ± 0.88 SE, *t* = −3.28, *p* = 0.002). In contrast, centered log body weight was not a significant predictor of spirocyst length (β = 0.33 ± 0.48 SE, *t* = 0.69, *p* = 0.496).

**Fig. 6:**
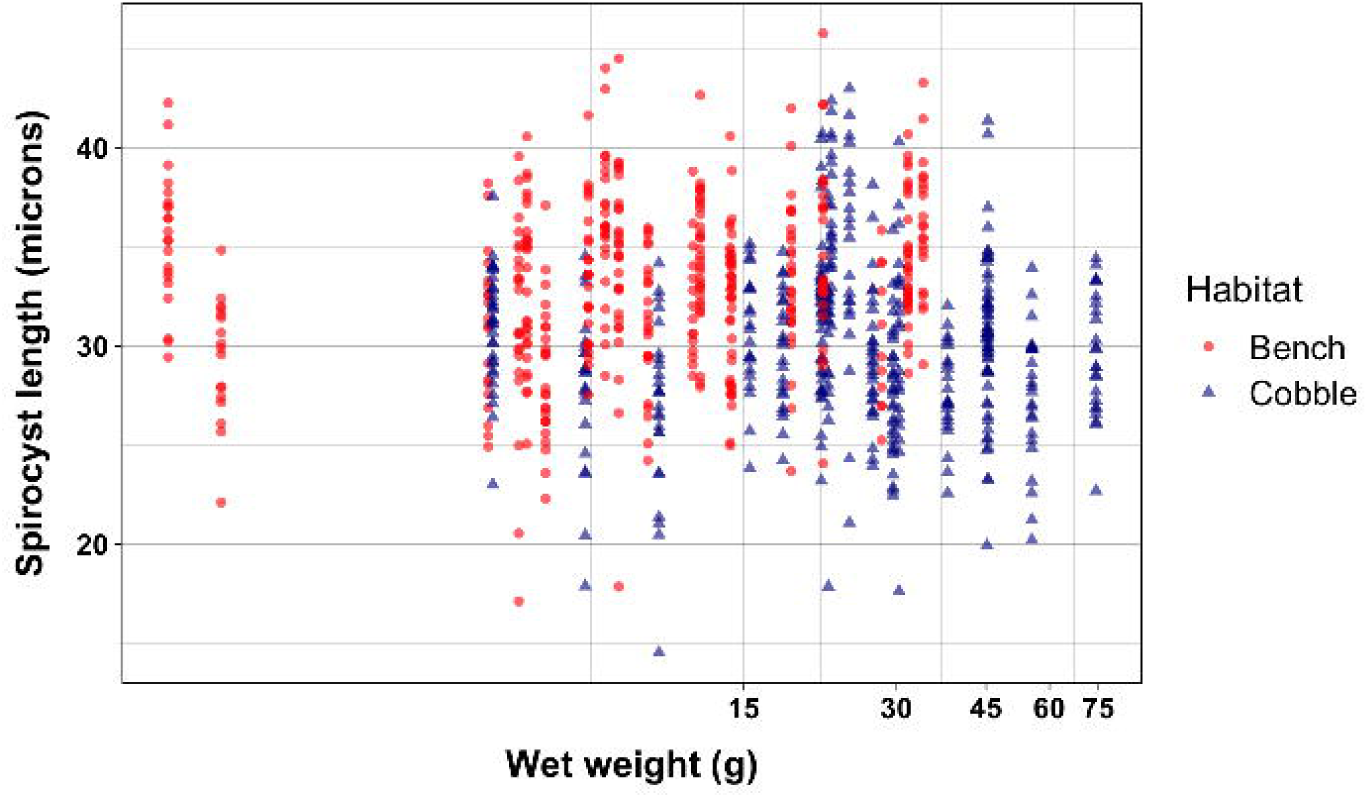
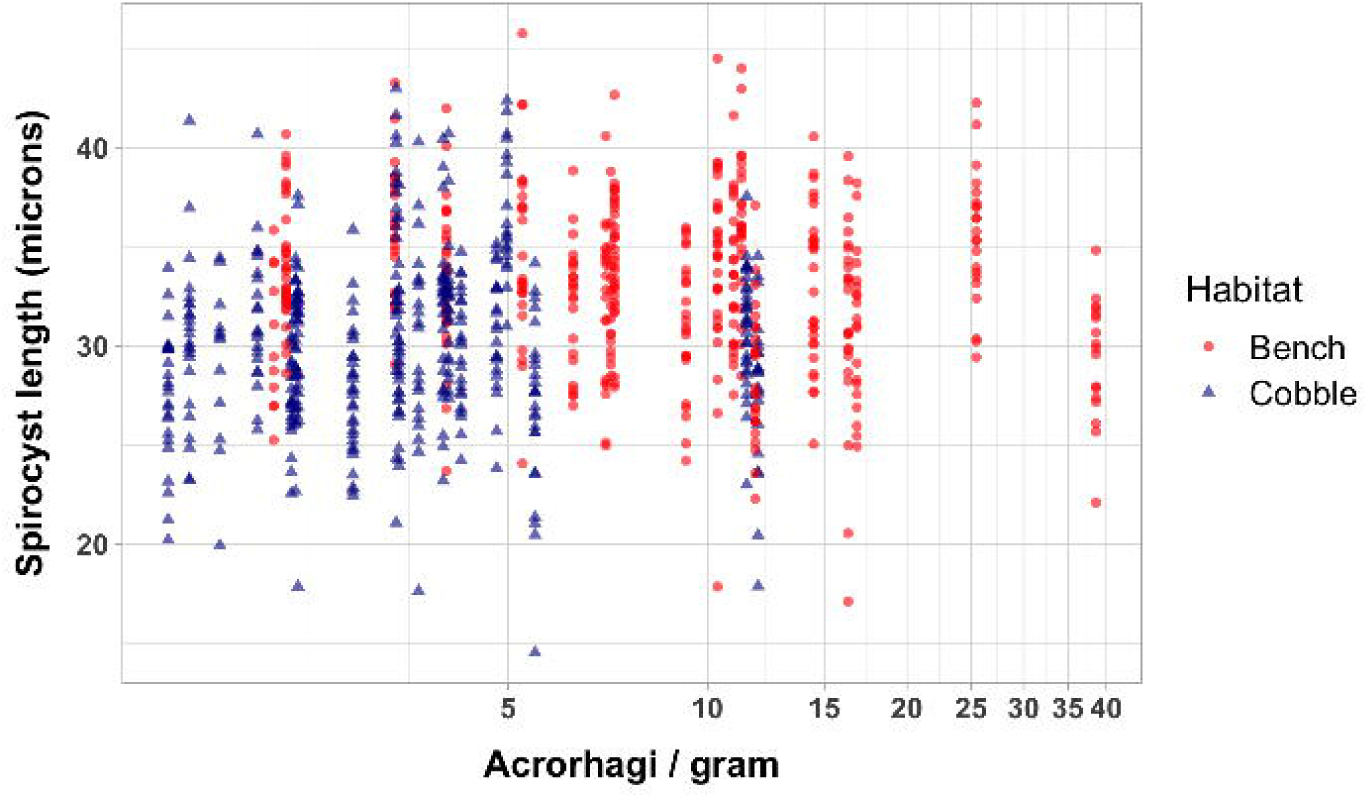
Spirocyst length (µm) is plotted against body weight (g) for individual anemones from bench (red circles) and cobble (blue triangles) habitats. Points represent individual spirocyst measurements. N= 19, N= 19

Spirocyst length differed significantly between habitat types after accounting for acrorhagial density (Fig. 7). In a linear mixed-effects model including habitat and centered log acrorhagi per gram as an additive effect, habitat had a significant effect on spirocyst length. Cobble anemones exhibited spirocysts approximately 2.84 µm shorter than bench anemones (β = −2.84 ± 0.92 SE, *t* = −3.09, *p* = 0.004). In contrast, centered log acrorhagial density was not a significant predictor of spirocyst length (β = −0.29 ± 0.59 SE, *t* = −0.49, *p* = 0.625).

**Fig. 7:**
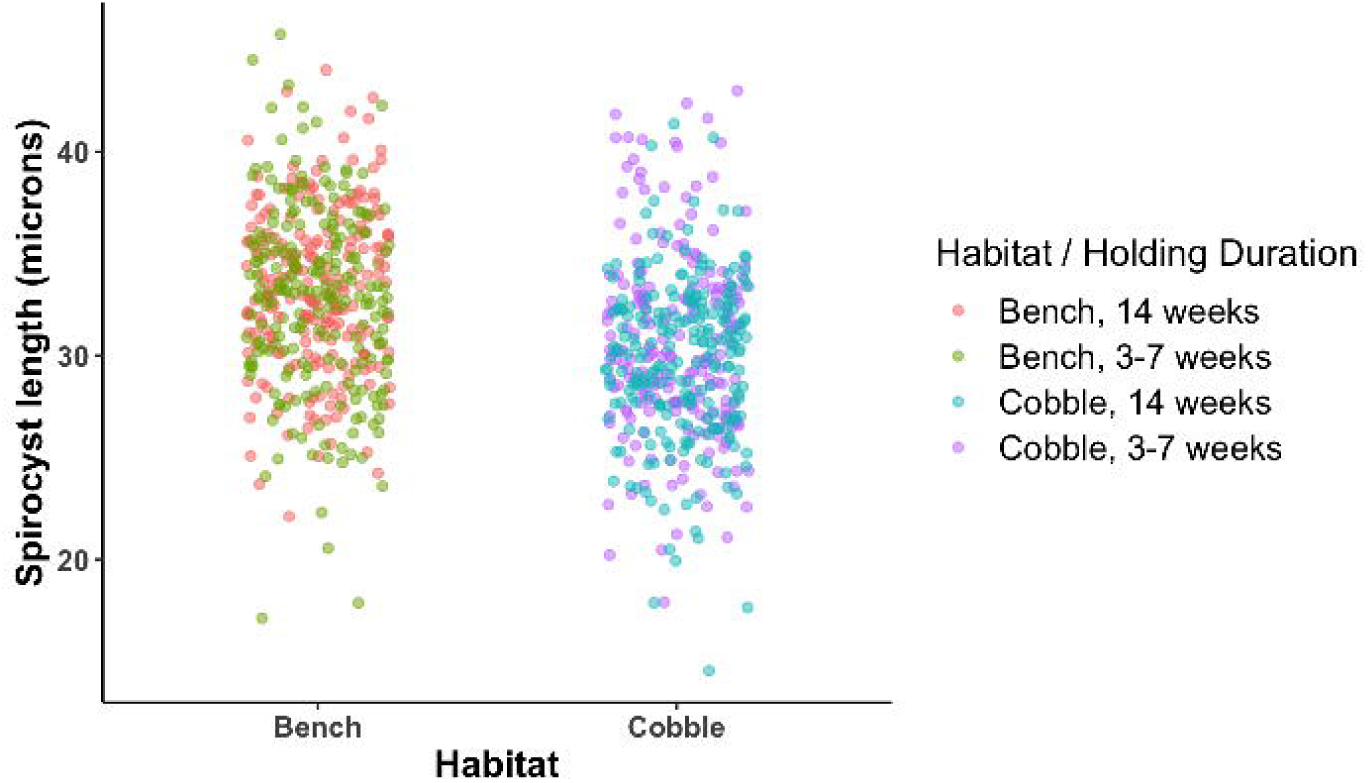
Spirocyst length (µm) is plotted against acrorhagi per gram for individual anemones from bench (red circles) and cobble (blue triangles) habitats. Points represent individual spirocyst measurements. N= 19, N= 19

There was no evidence of spirocyst length systematically differing between individuals processed earlier versus later within either habitat (suppl. Fig. 1), indicating that holding duration did not explain habitat-associated differences in spirocyst length. Numbers of acrorhagi do not change over short-term holding periods when anemones are not subjected to repeated contact with other individuals (Ayre and Grosberg, 2005; pers. obs. SVR).

## Discussion

The persistence of substantial intraspecific variation in weapon traits suggests that investment in weaponry may be regulated by variation in the local social environment. Here, we show evidence that weapon trait expression can covary with local environmental context. Directional differences in gross acrorhagial quantity were observed across habitat types, as well as at the level of the cnidom. These patterns suggest that variation in weapon morphology is not arbitrary, but may reflect plastic or selective mechanisms through which individuals modulate weapon development in response to local social conditions. In this way, our findings offer one potential explanation for the maintenance of morphological diversity in traits expected to be subject to strong ecological pressures. Additionally, we show that ecological pressures may simultaneously affect multiple levels of biological organization to influence weapon characteristics. To the best of our knowledge, this is the first study to simultaneously quantify intraspecific variation in both gross acrorhagial characteristics and acrorhagial cnidae.

### Acrorhagial anatomy and morphology

Variation in acrorhagial traits may reflect not only differences in investment, but also differences in the underlying anatomical architecture of the actiniid margin and body plan. We observe that the coelenteric space of acrorhagi in *A. sola* may communicate with either exocoelic or endocoelic compartments. This is in contrast to previous anatomical definitions which label acrorhagi as exclusively endocoelic structures (Daly, 2003). Yet, exocoelic acrorhagi have been observed (or implied) in other actinians (Den Hartog, 1987; Häussermann, 2004; Gomes *et al.,* 2012; V-Boada *et al*., 2026), as well as here for *A. sola*. Across acrorhagi-bearing actiniids, it seems that endocoelic acrorhagi are more prevalent than exocoelic acrorhagi (Daly, 2003; pers. obs. SVR).

The capacity to produce exocoelic acrorhagi appears to have significant allometric implications: forming an additional cycle of exocoelic acrorhagi allows an individual to effectively double its weaponry. Because different cycles of acrorhagi inflate in opposing directions (pers. obs. SVR), the effective surface area of the agonistic apparatus also increases, potentially intensifying the damage inflicted during contests. Species that can produce both endocoelic and exocoelic cycles may be adapted to especially intense competitive regimes. The production of exocoelic acrorhagi also appears to be body size-dependent, suggesting that evolving an increased body size may enable the elaboration of a secondary acrorhagial cycle. Given these biological implications, we propose that the presence of exocoelic acrorhagi is not only ecologically meaningful but may also be of taxonomic relevance.

### Acrorhagial scaling

As expected, the number of acrorhagi increased with anemone body size. This pattern is consistent with actinian growth, in which individuals add mesenteries as they increase in size, thereby expanding the number of endocoels and exocoels available to bear acrorhagi. However, mesenterial allometry alone cannot fully account for the scaling relationships observed here. Even relatively modest-sized anemones possessed numbers of mesenterial pairs approaching the modal condition, suggesting that acrorhagial production is driven by factors beyond increases in internal body plan complexity. Across habitats, and within each habitat, as anemones become larger, each additional gram contributes fewer new acrorhagi (negative allometry). This pattern contrasts with the positive allometry commonly reported for animal weapon size, which often exhibits disproportionate investment with increasing body size (Kodric-Brown *et al*., 2006). In this respect, acrorhagial development is constrained relative to other weapon systems.

With the exception of a few small bench individuals, our dataset contained limited representation of the lowest body-size classes. Mesenteries are added sequentially during growth, and early developmental stages may therefore exhibit disproportionately rapid increases in associated structures such as acrorhagi. In this way, the three smallest bench individuals may influence the estimated scaling exponent. More comprehensive sampling of early size classes would be required to determine whether scaling during early ontogeny deviates from the log–log relationship observed across the broader size range.

The maximum numbers of acrorhagi observed in the largest individuals (∼200) are comparable to the number of mesenteries present, indicating a structural upper bound on acrorhagial investment. Thus, acrorhagi differ from other structures of actinians: for example, they will be added at a slower rate than are tentacles, as a threshold in body size appears to need to be reached to begin adding to the exocoelic cycle. Such constraints fundamentally shape how weapon investment scales with body size in sea anemones. In addition to these size-dependent patterns, acrorhagial quantity showed evidence of differing between habitat types: anemones occupying the cobble habitat tended to possess fewer acrorhagi than did bench anemones of equivalent size. After accounting for body size, models indicated that cobble anemones possessed ∼20% fewer acrorhagi than bench anemones of equivalent mass. Statistical support for the habitat effect was modest and varied across model specifications, rendering strict significance sensitive to analytical approach. Nevertheless, the directional consistency of the effect indicates a potentially meaningful pattern that should be evaluated with greater statistical power.

### Spirocyst scaling

Spirocyst length did not demonstrate significant relationships with either anemone body weight or acrorhagial density. Yet, a consistently significant difference in mean spirocyst length (2.6–2.9 µm) was detected between habitats across model specifications, with cobble anemones possessing shorter spirocysts. The habitat effect persisted even when anemone weight and the density of acrorhagi were considered, indicating that differences in spirocyst length between habitats are not explained by variation in body mass or acrorhagi per unit body mass.

Our study adds to the evidence that the cnidom should not be treated as a fixed character (Fautin, 1988). The socio-environmental effects of variation in the cnidom may extend to other sea anemones with cnidae-laden weapons or defensive structures, such as catch tentacles or acontia (Kramer and Francis, 2004). We did not detect a size-scaling relationship in either habitat, or even across both habitats, in contrast to the significant scaling relationships that were found for *A. elegantissima* and *Anthopleura xanthogrammica* (Brandt, 1835) (Francis, 2004). Interestingly, larger-bodied anemones require thicker epithelial linings of the mesoglea, which may support larger cnidae (Francis, 2004). We show that environmental effects may potentially ‘override’ intrinsic physiology. While differences in cnidocyst size have been used to denote distinct morphotypes in several species of sea anemones on a scale on the order of kilometers (Acuña and Zamponi, 1997) and possibly across latitudinal gradients (Beneti *et al*., 2019), our study shows that such variation in cnidocyst morphology can occur over much smaller spatial scales (meters).

### Possible mechanisms

The close proximity of the two study habitats and the open recruitment of populations of many actinian species (Hunt and Ayre, 1989) suggest that true local adaptation is an unlikely explanation for the observed results. But this does not rule out other selective processes acting to differentially favor acrorhagial phenotypes, such as a balanced polymorphism (e.g., postsettlement selection). Such processes may maintain certain patterns of trait expression in anemones (Quicke *et al*., 1985) and other intertidal marine invertebrates (Sanford and Kelly, 2011).

Phenotypic plasticity may also play a significant role in driving these patterns of variation. Additional acrorhagi can be induced with repeated tentacular contact for the related *A. elegantissima* (Ayre and Grosberg, 2005), but the magnitude of these effects was inconsistent across individuals. Inducible agonistic structures are also common in other anthozoans such as scleractinians and corallimorphs (Chornesky, 1983; Miles, 1991). Plastic responses may extend beyond gross morphology to the cnidom as well (Langmead and Chadwick-Furman, 1999). Observations (pers. obs. SVR) that spatially isolated anemones nevertheless possess acrorhagi suggest that phenotypic plasticity alone does not fully explain acrorhagial production.

Although the most prominent difference between the two environments is conspecific density, these habitats differ in other respects (e.g., topography, trophic opportunities). Overall, the rocky intertidal zone is a heterogeneous landscape within which environmental parameters can vary across fine spatial scales (Dayton, 1971). Abiotic variables which may affect the cnidom are not well studied, although temperature differences were not found to have any significant effects on cnidae size in some cnidarians (Ling *et al*., 2025). We may also consider that dietary regimes of anemones in the rocky intertidal may differ on the order of meters (pers. obs. SVR). Gundlach and Watson (2019) showed that the cnidom of *Exaptasia diaphana* (Rapp, 1829) could be modulated in response to differential nutrient availability. However, even for pronounced experimental treatments, their effects on spirocyst volume were inconsistent, even though the tentacular cnidae directly function in capturing food. Lastly, relevant endogenous (life-history) factors of the anemones may also vary between habitats. The smaller size of bench anemones may facilitate movement, as has been shown for other actinian species (Bedgood *et al*., 2020), potentially increasing inter-individual encounter rates. This would still pertain to the competitive environment. As our study is associational, other confounding exogenous (environmental) or endogenous variables cannot be ruled out in explaining acrorhagial phenotypes. But given the proximity of the two habitats, as well as the specific function of the acrorhagi for use in intraspecific aggression, the difference in anemone density seems to be the most parsimonious explanation.

### Conclusion

Weapons are among the most conspicuous and well-studied traits in the animal kingdom, reflecting their central role in mediating competitive interactions. Yet, much of the empirical work on animal weapons has focused on their role in determining contest outcomes, rather than on the ecological contexts that shape their expression in the first place (Lane, 2018; Miller *et al*., 2026). In actiniid sea anemones, acrorhagi represent ecologically important traits that nevertheless exhibit substantial intraspecific morphological variation—variation that is likely to have functional consequences for competitive ability. Unlike many weapons that are typically studied, acrorhagi are not associated with mating success or sexual selection. Instead, they function in territorial encounters over space and access to resources. Because most work on animal weapons has focused on sexually selected systems, it remains unclear whether the ecological drivers of weapon expression, and the constraints under which they operate, generalize across these different functional contexts. Importantly, even within sexually selected systems, weapon expression can vary with ecological conditions; for example, investment may decline under high-density conditions subject to scramble competition, where costs of defending females exceed the benefits of weapon quality (Pomfret and Knell, 2008). These patterns suggest that the relationship between ecological context and weapon expression is not fixed, and may depend on both the functional role of the weapon and the environmental conditions in which it is deployed.

Population density represents a key component of ecological context that can influence weapon expression by mediating resource availability and competitive intensity. In some systems, weapon traits are constrained under high-density conditions due to resource limitation; for example, antler size in deer decreases with increasing population density, likely due to nutrient limitation (Schmidt *et al*., 2001). However, the biology of sea anemones may alter this relationship. As sessile organisms with flexible growth, anemones can adjust body size in response to resource conditions (Sebens, 1981), potentially allowing individuals to maintain investment in weapon traits even under crowding. This raises the possibility that the trade-offs constraining weapon expression in other taxa may be modified in this system.

Understanding the sources of intraspecific variation in such traits may also provide insight into their evolutionary dynamics. Weapons are often phylogenetically labile, with their form and function shaped by context-dependent costs and benefits that vary across environments (Miller *et al*., 2026). Even within sea anemones, acrorhagi exhibit notable diversity across clades, for example, in their association with endocoelic versus exocoelic compartments (Daly, 2003; this study), as well as differences in behavioral deployment during aggressive encounters, e.g., the reluctance to aggress in *A. xanthogrammica* (Bigger, 1980). These patterns suggest that both morphology and behavior of weapon traits may be evolutionarily flexible.

Finally, acrorhagi may be more appropriately viewed as a broader weapon system, in which gross morphological features (e.g., number and distribution of acrorhagi) and microscopic components (e.g., cnidae involved in venom delivery) together determine functional performance. Indeed, acrorhagi have been described as chemical weapons (Lane, 2018), insofar as the venoms contained within cnidae constitute the primary damage-inflicting agents. From this perspective, the present study focuses on two components of the weapon delivery system—morphological structure and cnidom characteristics—rather than the biochemical properties of the venom itself. While ecological pressures can nevertheless act on these ‘weapon-supportive’ traits (*sensu* Miller *et al*., 2026), a complete understanding of acrorhagial function in sea anemones may ultimately require integrating these components with variation in venom composition and potency.

## Data accessibility statement

The data that support the findings of this study are openly available on figshare at DOI: 10.6084/m9.figshare.32310912

## Acknowledgements

SVR conceptualized the study, collected and analyzed data, and primarily wrote the manuscript. JGS, CSK and ATB contributed to study design as well as data collection and analysis. THO provided intellectual guidance and supervision. All authors were involved in writing, revising, editing, and approving the final manuscript. Cristoph Pierre, Christian Orsini, Ryan Tang, Sophia Kaplan, Simren Gupta, Nicolas Vinas, Angela Ho, and Alyssa Prydz assisted in collection efforts. Marie Christodoulou assisted in data collection. We thank Armand Kuris, Rick Grosberg, and Lisbeth Francis for insightful discussions and encouragement. Partial funding was provided by the UCSB Associated Students Coastal Fund, awarded to SVR. All organisms were collected in accordance with permits S-223200008 and S-190950003-19142-001 issued by the California Department of Fish and Wildlife. This work was performed (in part) at the University of California Natural Reserve System (Coal Oil Point) Reserve DOI: (10.21973/N3Z07N).

