## Supplementary material for "Competitive environment predicts weaponry in an intertidal sea anemone"

### Structural anatomy of acrorhagi in *A. sola*

As stated in the text, acrorhagi are positioned at the distal edge of the body column (the margin) above the distal-most verrucae (tubercles on the body column). Acrorhagi (Fig. 2A,B) are specifically defined as bulbous protuberances containing dense clusters (or batteries) of holotrichous nematocysts (Daly, 2003) (but may also contain spirocysts or other nematocyst types, such as basitrichs). Like other external structures of sea anemones, acrorhagi communicate with the radial compartments formed by the partitioning of the coelenteron by pairs of sheet-like tissues termed mesenteries. These are termed endocoelic and exocoelic compartments (or simply, endocoels and exocoels), for those which lie within and between pairs of mesenteries, respectively (Fig. 2C). To date, acrorhagi have most frequently been characterized as exclusively endocoelic structures (Daly, 2003). For more detailed information on the marginal anatomy of actiniarian sea anemones, the reader is encouraged to consult Stephenson (1928) and Daly (2003).

### Mass correction function:

Eighteen individuals (n=9 per habitat) that were used in this study and held for an additional 16 weeks lost approximately between 12 and 55 % of their body mass. We modeled absolute mass loss ($\Delta M$) as a power function of the initial mass: $\Delta M = a_{habitat} M_0^{\ b}$ where $M_0$ is initial mass, $a_{habitat}$ is a habitat-specific rate constant describing the magnitude of mass loss, and $b$ is a shared scaling exponent describing how loss scales with body size. Fitted parameters were $a$ = 0.2574 for bench anemones, $a$ =

0.1454 for cobble anemones, and $b = 1.0718$ ($R^2 = 0.95$). These constants correspond to a 16-week holding period. The obtained scaling exponent ($b$) is similar to values in other models of anemone mass loss under starvation (Sebens, 1981).

The fitted parameters ($a$, $b$) were then used to estimate initial body masses ($M_0$ ) for all the anemones in the dataset, based on their measured masses at the time of processing ($M_t$) and known durations in holding ($t$, in weeks). To generalize the correction for individuals held for varying durations, we assumed mass loss scaled linearly with time $t$ over the intervals considered, such that: $M_t = M_0 - a_{habitat}(t/t_{\mathrm{ref}})M_0^{\,b}$ where $t_{\mathrm{ref}} = 16$; this first-order approximation avoids unsupported complexity given the absence of time-series measurements. Because this equation has no closed-form solution for $b \neq 1$, original mass was estimated numerically using fixed-point iteration. These corrected values represent the estimated body masses at the time of collection, accounting for size-dependent mass loss during holding.

**Supplementary figure legends:**

Suppl. Fig. 1: Spirocyst populations color-coded by habitat and holding duration. Red (Bench, 14 weeks), Green (Bench, 3−7 weeks), Blue (Cobble, 14 weeks), Purple (Cobble, 3−7 weeks)

**Suppl. Table 1: Counts of mesenterial pairs in selected individuals across both habitats.**

| Habitat | Individual weight (g) | Count of mesenterial pairs |
|---|---|---|
| Cobble | 30.1 | 101 |
| Cobble | 74.1 | 96 |
| Cobble | 36.0 | 95 |
| Cobble | 26.9 | 96 |
| Cobble | 29.1 | 94 |
| Cobble | 31.5 | 97 |
| Cobble | 19.8 | 96 |
| Cobble | 10.9 | 94 |
| Bench | 6.1 | 96 |
| Bench | 6.5 | 96 |
| Bench | 12.3 | 96 |
| Bench | 23.2 | 73 |
| Bench | 11.2 | 92 |

| | | |
|---|---|---|
| Bench | 13.1 | 96 |
| Bench | 37.0 | 92 |
